# Quantitative Superresolution Imaging of F-Actin in the Cell Body and Cytoskeletal Protrusions Using Phalloidin-Based Single-Molecule Labeling and Localization Microscopy

**DOI:** 10.1101/2024.03.04.583337

**Authors:** Hirushi Gunasekara, Thilini Perera, Chih-Jia Chao, Joshua Bruno, Badeia Saed, Jesse Anderson, Zongmin Zhao, Ying S. Hu

## Abstract

We present single-molecule labeling and localization microscopy (SMLLM) using dye-conjugated phalloidin to achieve enhanced superresolution imaging of filamentous actin (F-actin). We demonstrate that the intrinsic phalloidin dissociation enables SMLLM in an imaging buffer containing low concentrations of dye-conjugated phalloidin. We further show enhanced single-molecule labeling by chemically promoting phalloidin dissociation. Two benefits of phalloidin-based SMLLM are better preservation of cellular structures sensitive to mechanical and shear forces during standard sample preparation and more consistent F-actin quantification at the nanoscale. In a proof-of-concept study, we employed SMLLM to super-resolve F-actin structures in U2OS and dendritic cells (DCs) and demonstrate more consistent F-actin quantification in the cell body and structurally delicate cytoskeletal proportions, which we termed membrane fibers, of DCs compared to direct stochastic optical reconstruction microscopy (*d*STORM). Using DC2.4 mouse dendritic cells as the model system, we show F-actin redistribution from podosomes to actin filaments and altered prevalence of F-actin-associated membrane fibers on the culture glass surface after lipopolysaccharide exposure. While our work demonstrates SMLLM for F-actin, the concept opens new possibilities for protein-specific single-molecule labeling and localization in the same step using commercially available reagents.

## INTRODUCTION

The actin cytoskeleton consists of distinct filamentous actin (F-actin) arrangements, including stress fibers, cortical actin, lamellipodia, and filopodia, which regulate cellular dynamics, provide mechanical support, maintain the cell shape, and modulate cell migration. Fluorescent labeling of F-actin using dye-conjugated phalloidin has enabled extensive superresolution studies ^1–7^ using single-molecule localization microscopy (SMLM). ^8–11^

F-actin is also known to regulate the morphology and function of cytoskeletal protrusions. Recently, actin-enriched elongated membrane structures have been shown to play increasingly important roles in the progression of various pathologies at the nanoscale, including viral and bacterial infections, ^12–15^ cancer, ^16–18^ and neurodegenerative diseases. ^19^ In addition, these membrane structures carry out physiological functions, such as mitochondrial homeostasis, ^20^ immune surveillance, ^21,22^ endocytosis, ^23^ and exosome release. ^24^ In particular, their involvement in intercellular communications provides new opportunities for drug delivery and therapeutic intervention. ^16,25,26^ Electron microscopy (EM) ^16,27–29^ and correlative light-electron microscopy (CLEM) techniques have been used to study membrane fibers and the presence of F-actin within these fibers. ^30^ Extending SMLM techniques to study morphology and F-actin distribution in these delicate cellular structures requires tailored sample preservation methods. ^26^ The integrating exchangeable single-molecule localization (IRIS) of F-actin and points accumulation for imaging in nanoscale topography (PAINT) using transient interactions of fluorophore-conjugated Lifeact ^31^ present alternative strategies to achieve superresolution imaging of F-actin with higher labeling densities. ^32,33^ However, the binding of actin probes to F-actin depends on the precise biochemical interactions dictated by the F-actin nano-architecture^34^, and Lifeact may not efficiently label fragile cellular structures, such as filopodia in mesenchymal cells^35^ and protein-bound actin in mouse striatal neuron-derived STHdh cells. ^36^ Therefore, a gentle labeling strategy with an actin probe widely used on membrane fibers, such as phalloidin, will extend SMLM to study fragile cellular structures.

Here, we present a single-molecule labeling and localization microscopy (SMLLM) technique using dye-conjugated phalloidin (phalloidin-based SMLLM). We show that dye-conjugated phalloidin displays moderate dissociation from the fixed cell sample. By promoting the dissociation using a chaotropic agent, we achieved superresolution imaging and molecular quantification of F-actin in U2OS and immortalized (DC2.4) and bone marrow derived primary mouse dendritic cells (BMDC). We further demonstrate superresolution images of SMLLM revealing the spatial redistribution of F-actin and diminished F-actin-associated membrane fiber networks of DC2.4 cells after exposure to lipopolysaccharide. Thus, phalloidin-based SMLLM provides a simple and versatile technique for quantitative superresolution imaging of F-actin.

## RESULTS AND DISCUSSION

### Single-molecule labeling and localization microscopy using dye-conjugated phalloidin achieve superresolution imaging of F-actin

We selected the widely used Alexa Fluor^TM^ 647 conjugated phalloidin (phalloidin-AF647) as the dye-conjugated phalloidin for our study. As phalloidin-dye conjugates are reported to show intrinsic faster dissociation from F-actin,^33,37^ we first characterized the loss of phalloidin-AF647 staining in DPBS. We stained DC2.4 cells with phalloidin-AF647 and monitored the fluorescent intensity over time. Consistent with previous reports, the fluorescence signal from phalloidin-staining of DC2.4 dropped by approximately 40% - 50% within 40 min in DPBS (**Figure 1a-d**).^33,37^ The consecutive images of the same cell exhibited no significant change in the fluorescence intensity, indicating a minimal impact from photobleaching (**Figure 1a**). Although F-actin-rich podosome structures remained distinct after 40 min, the membrane structures were barely visible (**Figure 1a-d**). We further characterized the fluorescence signal loss in the STORM buffer, which is used in direct stochastic optical reconstruction microscopy (*d*STORM) superresolution imaging of phalloidin-labeled F-actin. The fluorescence signal was reduced even more in STORM buffer by approximately 70% - 85% within 40 min (**Figure 1e-h**). STORM buffer contains 2-mercaptoethanol to facilitate the photoswitching of AF-647. Thus, the additional fluorescence signal loss in the STORM buffer may be due to 2-mercaptoethanol attacking the pi system of AF647.^38^ We found this to be valid as the fluorescence signal after 40 min could be partially recovered by stimulating the sample with a 405 nm laser.

**Figure 1.**
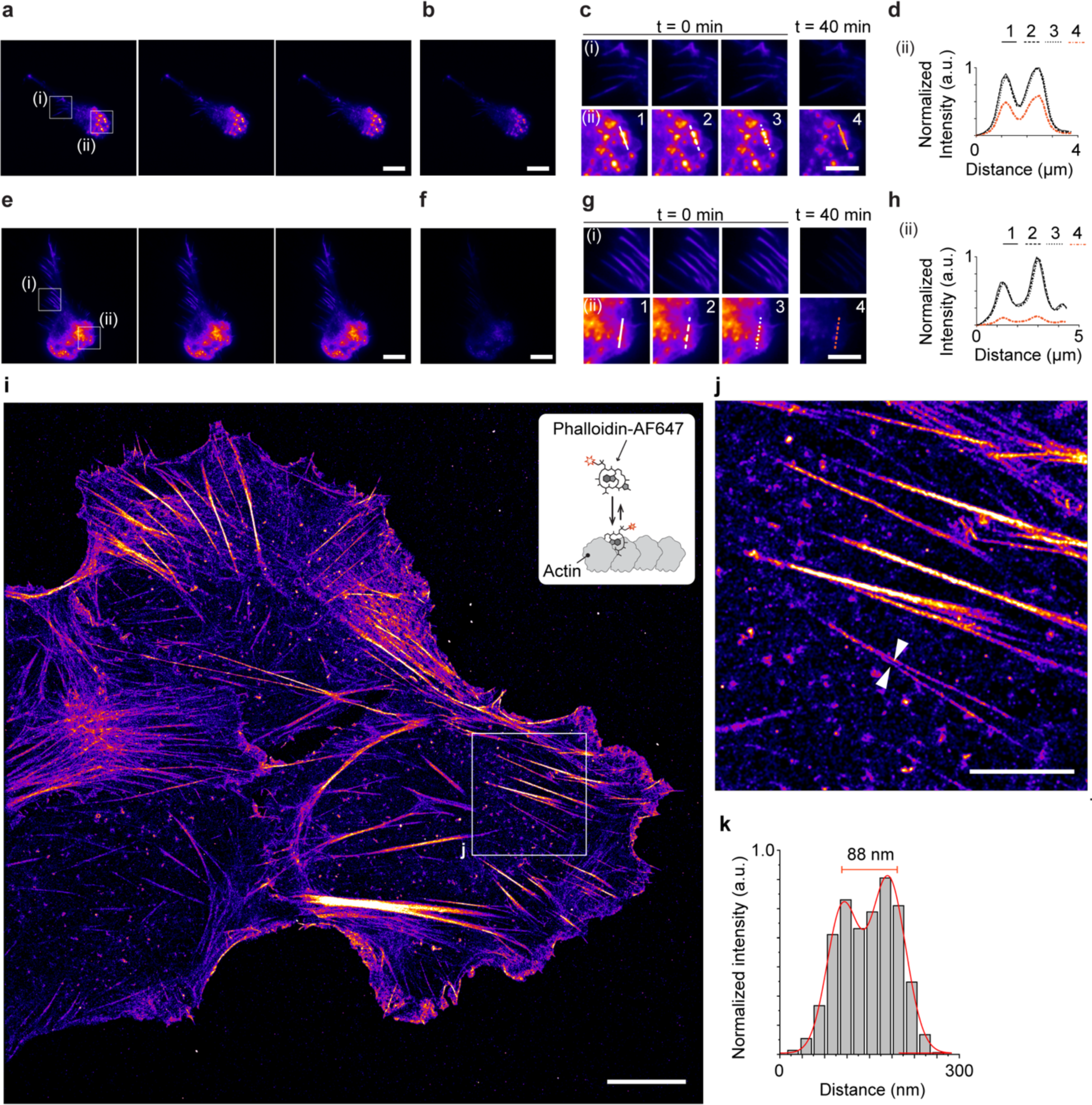
Intrinsic dissociation and single-molecule labeling with phalloidin-AF647 enabled superresolution imaging of F-actin. **a.** Three consecutively acquired fluorescence images of the same DC2.4 cell stained with phalloidin-AF647 incubating in 1xDPBS buffer. **b.** A fluorescence image of the DC2.4 cells shown in panel a, acquired after 40 min. Image contrast is adjusted to that of panel a. **c.** Zoomed in views of the boxed regions in panel a, across panels a and b. **d.** Corresponding cross-sectional intensity profile along the lines indicated in panels c (ii). **e.** Three consecutively acquired fluorescence images of the same DC2.4 cell stained with phalloidin-AF647. Images were acquired in the STORM buffer immediately after adding the buffer. **f.** A fluorescence image of the DC2.4 cells shown in panel e, acquired after 40 min. Image contrast is adjusted to that of panel e. **g.** Zoomed-in views of the boxed regions in panel e, across panels e and f. **h.** Corresponding cross-sectional intensity profile along the lines indicated in panels g (ii). **i.** A representative SMLLM superresolution image reconstructed from a 30,000-frame acquisition with phalloidin-AF647 on a U2OS cell (right) and a schematic illustration of single-molecule labeling of F-actin with phalloidin-AF647 (inset). **j.** Zoomed-in view of the boxed region in panel i. **k.** Gaussian-fitted cross-sectional profile across the F-actin fibers indicated in panel j. Scale bars: 10 μm (**a**, **b**, **e**, **f** and **i**) and 5 μm (**c**, **g** and **j**).

Phalloidin is a rigid bicyclic heptapeptide small molecule actin probe with specificity and affinity to F-actin in cells.^39^ Given the comparable size of phalloidin (790 Da) to the conjugated small molecule dyes (AF-647 1160 Da), the conjugated dye influences its interaction with the F-actin. Early reports on phalloidin conjugates report a ‘relative affinity’ to phalloidin conjugates compared to unconjugated phalloidin. ^40–42^ Cationic fluorophores like rhodamine may introduce extra electrostatic attraction (relative affinity of 2), while anionic fluorophores like fluorescein may repel from F-actin (relative affinity of 0.1). ^40^ By the same token, electrostatic repulsions from the anionic AF647 ^43^ might explain the observed intrinsic dissociation of the phalloidin-AF-647.

Next, we utilized U2OS cells as our model system to demonstrate single-molecule labeling and localization microscopy (SMLLM) of F-actin. Our approach involves reducing the concentration of phalloidin to decrease its binding rate, enabling us to observe and record individual phalloidin binding events in a single-molecule imaging setup. ^44^ In a proof-of-concept experiment, we performed SMLLM on fixed U2OS cells with phalloidin-AF647 under a highly inclined and laminated optical sheet (HILO) illumination. We introduced a non-illuminating interval of 2 s between consecutive image frames to ensure adequate detection of single-molecule events without significant spatial overlap. ^44^ **Figure 1i** shows a superresolution image reconstructed from a 30,000-frame movie. While single-molecule labeling of phalloidin-AF647 successfully resolved thick F-actin stress fibers (**Figure 1i**), we observed inadequate representation of the thin actin filaments (**Figure 1j**). This limitation is likely due to the slower sampling rates resulting from the relatively slow interaction kinetics between phalloidin and F-actin. To improve the sampling rate, we destabilized the phalloidin-AF647-F-actin interaction.

### Promoting phalloidin dissociation through chaotropic perturbation enhances F-actin labeling density and quantification consistency of phalloidin-based SMLLM

Phalloidin interacts with F-actin through hydrophobic interactions. ^34,45,46^ We have previously demonstrated the capability of chaotropic perturbation of hydrophobic interactions to promote dissociation rates between antibody fragments and their polypeptide ligands to achieve superresolution molecular census. ^47^ Here, we employed the chaotropic salt potassium thiocyanate (KSCN) to further enhance the dissociation of phalloidin from F-actin (**Figure 2a**, left). We incubated fixed U2OS cells with phalloidin-AF647 in an imaging buffer supplemented with 300 mM KSCN (methods). **Figure 2a** (right) shows a corresponding superresolution image reconstructed from a 30,000-frame movie. The chaotropic perturbation enhanced the sampling rate, enabling us to resolve stress fibers (**Figure 2b**) and thin F-actin filaments (**Figure 2c**). **Figure 2d** demonstrates the superresolution achieved by chaotropic perturbation-enhanced single-molecule labeling with phalloidin-AF647, resolving the adjacent F-actin fibers by approximately 75 nm at the point indicated in **Figure 2b**. To colocalize the superresolution images and validate that the phalloidin-based SMLLM localizations align with those obtained using the standard F-actin SMLM technique, *d*STORM, we performed single-molecule labeling with phalloidin-AF647 on the same cell after performing *d*STORM imaging (**Figure 2e-g**). The overlaid image demonstrates the successful colocalization of the phalloidin-based SMLLM image with the *d*STORM image, represented as white regions and along the thin fibers (**Figure 2g**, arrowheads). Thus, single-molecule labeling of F-actin recapitulated the F-actin structures resolved by *d*STORM imaging, showcasing the precise capture of F-actin.

**Figure 2.**
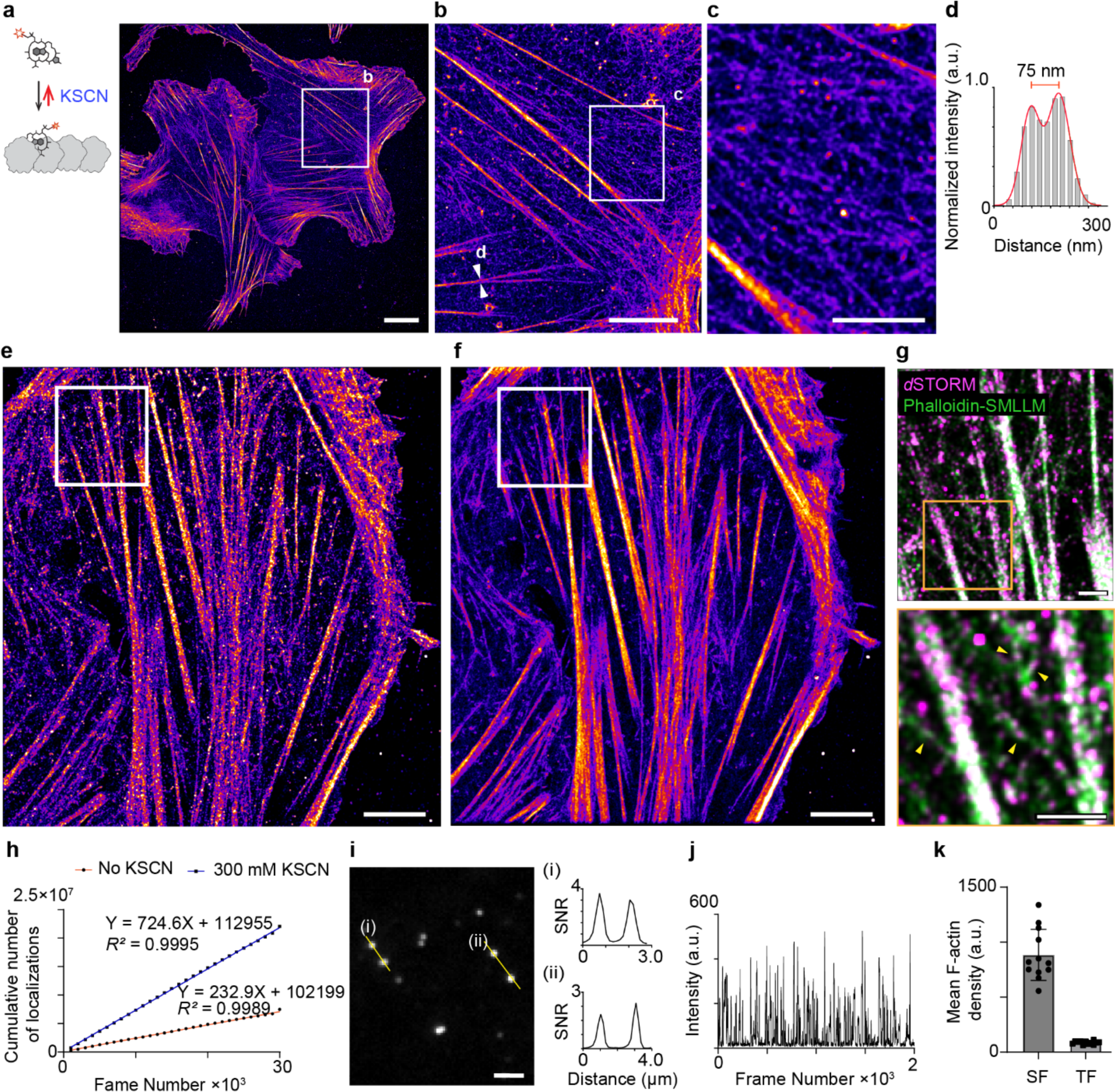
Chaotropic perturbation enhances F-actin labeling consistency and quantification using SMLLM. **a.** Schematic illustration of chaotrope (KSCN) enhanced single-molecule labeling of F-actin (left) and a representative superresolution image reconstructed from a 30,000-frame acquisition with phalloidin-AF647 in the presence of 300 mM KSCN on a U2OS cell (right). **b.** Zoomed-in view of the boxed region in panel a. **c.** Zoomed-in view of the boxed region in panel b. **d.** Gaussian-fitted cross-sectional profile across the F-actin fibers indicated in panel b. **e.** *d*STORM image of a phalloidin-stained U2OS cell. **f.** The phalloidin-based SMLLM image of the U2OS cell shown in panel e immediately after the *d*STORM acquisition. **g.** Merged view of the boxed region shown in panels e and f (top). The Zoomed-in view of the yellow boxed region indicated (bottom). Yellow arrowheads point to the thin actin fibers. **h.** Linear regression fit of the cumulative number of events *per* 1,000 frames in the presence (squares, blue fit) and absence (circles, orange fit) of 300 mM KSCN. **i.** Representative single-molecule localization events from a phalloidin-based SMLLM acquisition in the presence of 300 mM KSCN. The cross-sectional intensity profile across the indicated yellow lines shows the SNR. **j.** Representative time trace of single-molecule events detected with phalloidin-AF647 on a fixed U2OS cell in 300 mM KSCN. **k.** Quantitative comparison of the relative F-actin densities in stress fibers (SF) and thin fibers (TF). Scale bars: 10 μm (**a**), 5 μm (**b**, **e**, and **f**), 2 μm (**c** and **i**), and 1 μm (**g**).

To evaluate the sampling rates quantitatively, we fitted the cumulative number of single-molecule events *per* 1,000 frames to a linear regression model. The slope of the plot depicted in **Figure 2h** corresponds to the average number of single-molecule events detected *per* frame. Notably, the presence of KSCN in the single-molecule labeling image acquisitions resulted in an approximately 3-fold increase in the sampling rate. Moreover, the progressive and linear accumulation of localizations suggests that F-actin was continuously probed throughout the image acquisition, maintaining a constant sampling rate over an extended period of 16 h. **Figure 2i** shows the individual phalloidin-F-actin interactions under the HILO illumination, manifested as distinct single-molecule events with an average signal-to-noise ratio (SNR) of 4. **Figure 2j** shows a single-molecule intensity track exhibiting single-molecule binding events detected over 2,000 frames acquired in the presence of KSCN. The distinct intensity profiles in **Figure 2j** represent phalloidin-AF647 probing F-actin in the cell continuously. Since phalloidin-based SMLLM directly probes F-actin, the intensity of the 2D probability histogram represents the local F-actin population densities. We evaluated the F-actin densities across thick stress fibers and thin fibers of the cell shown in **Figure 2f** by taking mean intensity profiles (Figure S1). **Figure 2k** shows that the F-actin density of stress fibers of the U2OS cell analyzed is approximately 10 times higher than that of thin fibers.

### Phalloidin-based SMLLM reveals F-actin-associated membrane fiber structures in immortalized and primary dendritic cells

Next, we investigated the nanoscale F-actin arrangements in membrane structures of DC2.4 mouse dendritic cells. Phalloidin-based SMLLM of DC2.4 cells revealed actin polarization at the leading edge of the DC2.4 cell, manifested as F-actin-rich podosomes. Furthermore, phalloidin-based SMLLM illustrated that these membrane fibers were relatively rich in actin, similar to podosomes (**Figure 3a** and **b**). In contrast, the local actin densities in the rest of the cell body appeared much lower. In addition to resolving F-actin-associated membrane structures near the trailing edge of the cell, phalloidin-based SMLLM also revealed distal membrane fibers with low actin contents (Figure S2). We confirmed the low actin content observed with phalloidin single-molecule labeling using SiR-actin labeling on live DC2.4 cells. Figure S3 presents the fluorescence image of SiR-actin-labeled live DC2.4 cells and representative line profiles through a DC2.4 cell and membrane fibers for comparative analysis. Notably, the intensity of the membrane fibers was at least approximately 500 times lower than the average intensity of the cell body (Figure S3b). We further determined the thickness of the F-actin in distal membrane fibers from two independent experiments to be 52 ± 6 nm (*n = 82*) (Figure S2c-d). The measured actin fiber thickness was smaller than our previously measured membrane fiber thickness of 150 nm, ^21^ suggesting additional luminal space to accommodate actin-associated motor proteins for cargo transfer. ^48^

**Figure 3.**
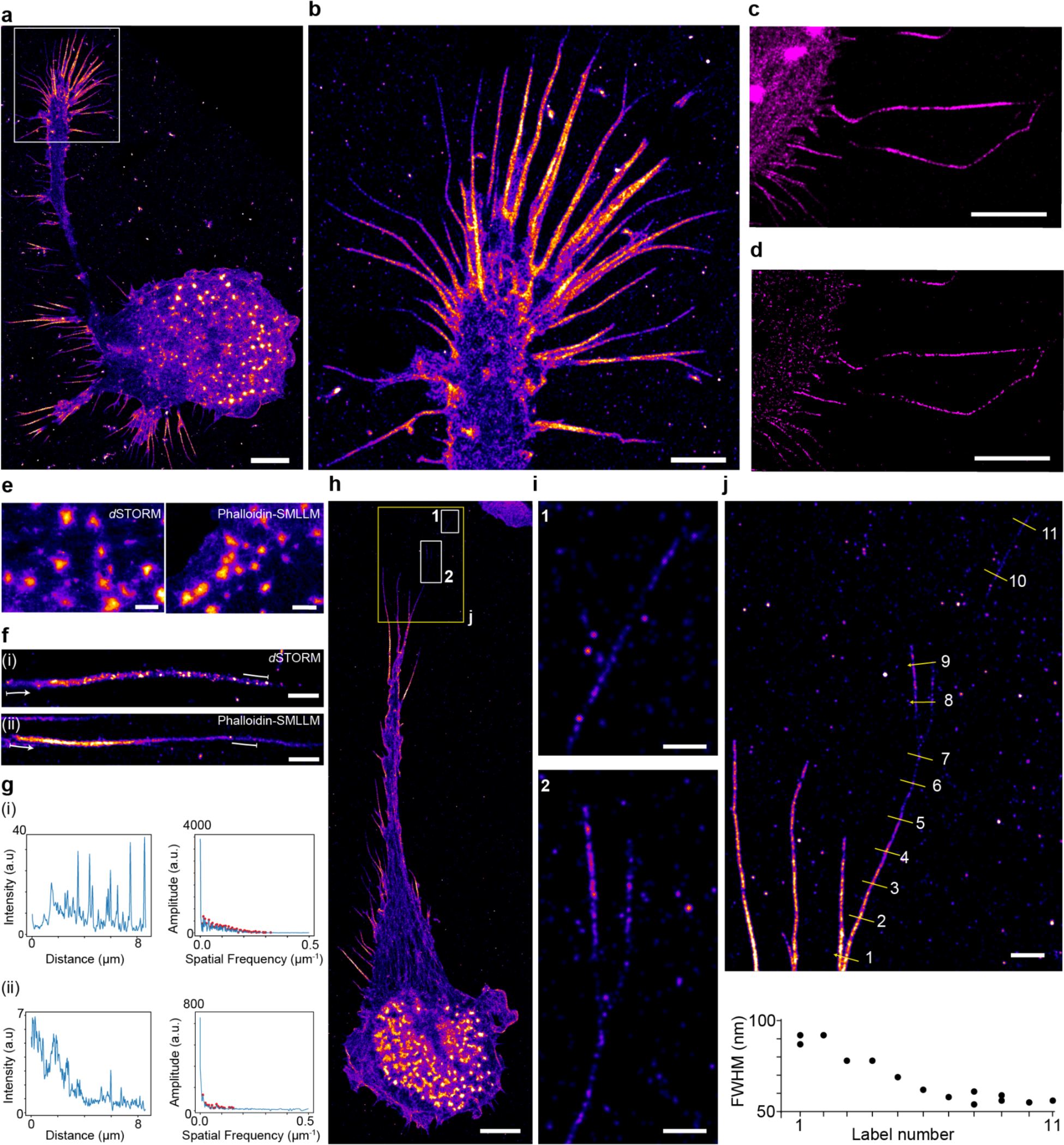
Phalloidin-based SMLLM superresolution imaging of F-actin in podosomes and membrane fibers of dendritic cells. **a.** A whole-cell phalloidin-based SMLLM image of a DC2.4 cell. **b.** Zoomed-in view of the boxed region in a. **c.** Phalloidin-based SMLLM superresolution image of DC2.4 cell (before washing steps). **d.** Structures lost during the washing step. The image was obtained by subtracting the phalloidin-based SMLLM image after washing from before washing (in panel c). **e.** Representative podosome structures from a *d*STORM experiment (left) and a phalloidin-based SMLLM experiment (bottom). **f.** Representative membrane fibers reconstructed from a *d*STORM experiment (top). Representative membrane fibers from a phalloidin-based SMLLM experiment (bottom). **g.** Evaluation of the labeling consistency along membrane fibers reconstructed from *d*STORM (i, top) and phalloidin-based SMLLM (ii, bottom). Cross-sectional intensity profile along the membrane fiber in panel f(i) (top left) and the corresponding FFT plot (top right). The red circles highlight detected frequency components. Cross-sectional intensity profile along the indicated membrane fiber in panel f(ii) (bottom left) and the corresponding FFT plot (bottom right). The red circles highlight detected frequency components. **h.** A whole-cell phalloidin-based SMLLM image of a primary bone marrow derived dendritic cell (BMDC) from mice. **i.** Contrast adjusted zoomed-in view of the numbered boxed regions indicated in h. **j.** Variation of the fiber thickness moving outward from the cell body. The contrast-adjusted zoomed-in region is indicated in panel j (top). The fiber thickness was measured at FWHM along the indicated points in the image (bottom). Scale bars: 5 μm (**a**), 2 μm (**b**), 3 μm (**c** and **d**), 1 μm (**e**, **f** and **j**), 10 μm (**h**), and 0.5 μm (**i**).

Membrane fiber structures are fragile and can be readily damaged during solution exchange steps. Standard superresolution experiments include a phalloidin-staining step, and the stained samples subsequently require multiple washing steps to remove excess fluorescent phalloidin before adding the corresponding imaging buffer. To evaluate the impact of washing steps on the membrane fiber structures of DC2.4, we first performed phalloidin-based SMLLM on DC2.4 cells. To imitate the conventional washing steps, we washed the sample three times immediately following single-molecule labeling acquisition. We subsequently re-imaged the sample with phalloidin-based SMLLM. **Figure 3c** shows the superresolution image of the DC2.4 cell before the washing steps. We subtracted the superresolution image after the washing steps from the superresolution image before the washing steps such that the resultant image would show structures present before the washing steps but absent after the washing steps (**Figure 3d**). **Figure 3d** shows the resulting image showing the membrane fiber structures, suggesting these structures were vulnerable to shear forces during the washing steps. Therefore, phalloidin-based SMLLM presents a gentle labeling strategy for superresolution imaging of fragile structures. While we demonstrate the influence of washing steps on membrane fiber structures of DC2.4, the influence of washing steps on the membrane fiber structures could vary across cell types, depending on the precise F-actin nanoarchitecture and mechanical strength of the actin-associated membrane fibers.

Phalloidin-based SMLLM demonstrated high labeling density on DC2.4 membrane fiber structures. We evaluated the labeling consistency and density between *d*STORM and phalloidin-based SMLLM on podosomes and membrane-fiber structures. Podosomes, the F-actin-rich mechanosensory apparatus of the DC, have been extensively characterized using superresolution methods, including *d*STORM. ^2,49^ Phalloidin-based SMLLM resolved podosome structures *on par* with *d*STORM (**Figure 3e**). In contrast, phalloidin-based SMLLM displayed a higher labeling consistency than *d*STORM on membrane fibers (**Figure 3f-g and** Figure S4). We evaluated the labeling consistency of membrane fibers resolved by *d*STORM and phalloidin-based SMLLM in the frequency space by obtaining the fast Fourier transform (FFT) of the cross-sectional intensity profiles (**Figure 3g** and Figure S5). The FFT analysis shows that the spatial frequency components extend further into high frequencies for *d*STORM reconstructions. In contrast, phalloidin single-molecule labeling exhibits fewer spatial frequency components (**Figure 3g** and Figure S5a *vs.* b). The segregated appearance of *d*STORM reconstructions could be attributed to either a loss of labeling during data collection from photobleaching and dissociation ^33,37^ or blinking artifacts. ^50^ Phalloidin-based SMLLM is immune to these artifacts.

Membrane fiber structures were also observed in primary dendritic cells. **Figure 3h** shows phalloidin-based SMLLM images of a primary bone marrow-derived dendritic cell (BMDC) isolated from BALB/c mice (methods). We further observed thin actin filaments extending away from the cell body with a relatively low actin density compared to the cellular actin structures (**Figure 3i-j**). **Figure 3i** and **j** display contrast-adjusted, zoomed-in views of these structures. In addition to a reduction in actin density, the fiber thickness decreased, moving away from the cell body. We characterized the thickness of the F-actin in the membrane fibers by generating cross-sectional profiles across the fiber (numbered in **Figure 3j**, top). The full-width-at-half-maximum (FWHM) measurements showed that the individual fibers originated with a thickness of approximately 100 nm, gradually decreased in thickness, and became consistent at approximately 55 nm (**Figure 3j**, bottom). Our observations of DC2.4 cells and BMDCs suggest that actin-associated membrane fibers are characteristic of immature dendritic cells.

### Phalloidin-based SMLLM reveals F-actin rearrangements in DC2.4 dendritic cells after LPS exposure

LPS activates pattern recognition receptors (TLR4) to trigger DC cell maturation, ^51^ and treatment of DC with lipopolysaccharide (LPS) induces cytoskeletal rearrangements that facilitate DC migration to the lymph nodes (**Figure 4a**). ^52,53^ To visualize the nanoscale F-actin rearrangement in DC2.4, we stimulated DC2.4 cells with LPS for 21 h ^53,54^ and performed phalloidin-based SMLLM. **Figure 4b, c**, and Figure S6 show representative single-molecule labeling images of immature DC2.4 (iDC2.4, without LPS treatment) and LPS-treated DC2.4 (LPS-DC2.4), respectively. Consistent with the literature, ^52,53,55,56^ LPS-DC2.4 cells showed loss of podosome structures, no apparent actin polarization, and distinct dendrite formation compared with the iDC2.4 cells (**Figure 4c** *vs.* **Figure 4b**). Phalloidin-based SMLLM also captured membrane fibers in both conditions (**Figure 4d** and **e**). Furthermore, in comparison to the fine mesh-like actin cytoskeleton of the iDC2.4, the actin cytoskeleton of the LPS-DC2.4 cells appeared to be much more defined, shaping the extending dendrite morphology (**Figure 4g** *vs.* **Figure 4f** and Figure S7). The observed rearrangement of the actin cytoskeleton, along with the generation of more dendrites, the decrease in the number and rearrangement of membrane fibers in mature DCs likely aligns with their functional shift from antigen survey to antigen presentation.^55^

**Figure 4.**
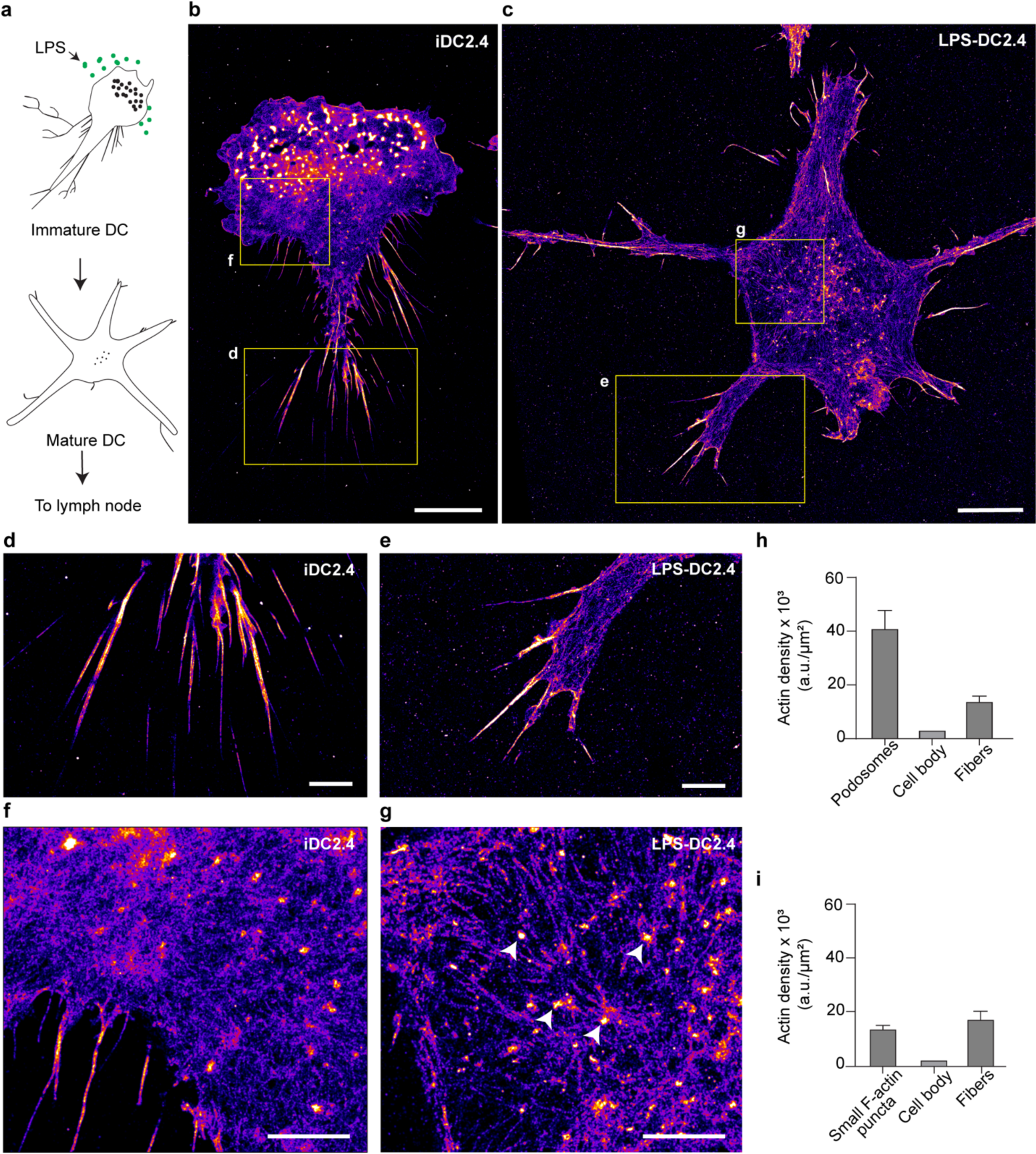
Phalloidin-based SMLLM reveals nanoscale F-actin rearrangements upon LPS treatment of DC2.4 cells. **a.** Schematic illustration of the altered cellular morphology in response to LPS activation. **b.** Phalloidin-based SMLLM of an iDC2.4 cell. **c.** Phalloidin-based SMLLM image of an LPS-DC2.4 cell. **d.** Zoomed-in view of the indicated region in panel b showing membrane fibers in iDC2.4. **e.** Zoomed-in view of the indicated region in panel b showing membrane fibers in LPS-DC2.4. **f.** Zoomed-in view of the indicated region in panel b showing cytoskeletal actin arrangement of iDC2.4. **g.** Zoomed-in view of the indicated region in panel c showing cytoskeletal actin arrangement of LPS-DC2.4. **h.** Distribution of actin density across the podosomes, cell body, and membrane fiber structures in iDC2.4. Error bars represent standard deviation. **i.** Distribution of actin density across the actin-dense regions, the cell body, and membrane fiber structures in LPS-DC2.4. Error bars represent standard deviation. Scale bars: 10 μm (**b** and **c**) and 3 μm (**d-g**).

Phalloidin-based SMLLM also enables quantitative assessment of local actin concentrations within a DC. Based on the intensity of the superresolution image, we conducted a global analysis of the actin densities of the cells depicted in **Figure 4b** and **c**. Our analysis revealed three distinct actin populations in the iDC2.4 and LPS-DC2.4 cells: subcellular actin-dense regions (podosomes in iDC2.4 in LPS-DC2.4), cell body, and membrane fibers (Figure S8 and **Figure 4h-i**). To facilitate comparison, we assessed the density of actin-rich regions and membrane fibers compared to that of the cell body. Of note is that the local actin density in the cell body and membrane fibers remained relatively unchanged before and after LPS exposure (**Figure 4h** *vs.* **i**), which likely reflects the enhanced cell-to-cell quantification consistency of SMLLM. Furthermore, **Figure 4g** displays the actin densities for iDC2.4 through podosomes, cytoskeleton, and membrane fibers. In the cell shown, the actin density in the podosomes was much higher (*c.a.,* 12-fold) compared to that of the cell body. In addition, proximal membrane fibers also displayed a higher actin density (*c.a.,* 4-fold) relative to the cell body. In contrast, intracellular actin appeared to be redistributed into more pronounced cytoskeletal structures in LPS-DC2.4. However, interestingly, LPS-DC2.4 cells displayed some subcellular actin-rich regions (*c.a.,* 6-fold higher) compared with the cell body. These actin-rich regions were distributed in the cell body and along the cell periphery, as evident in **Figure 4g** (arrowheads) and Figure S8, respectively. The latter observation aligns with existing literature that cortical actin stiffens upon maturation to support effective antigen presentation to T-cells.^57^ While membrane fibers were much less prevalent in LPS-DC2.4, the local actin densities remained higher (**Figure 4i**, *c.a.,* 7-fold) than the cell body. In both cases of iDC2.4 and LPS-DC2.4, the relative actin density of the membrane fibers was higher than that of the cell body, indicative of the heightened activity of these structures. These results suggest spatial redistributions of F-actin into nanoarchitecture to accommodate DC’s physiological functions, *i.e.,* from antigen sampling and sensing to cell migration and antigen presentation with LPS treatment.

In summary, we present a single-molecule labeling strategy for superresolution imaging of F-actin in fragile actin-associated membrane fiber structures using phalloidin-AF647. We have successfully showcased the performance of phalloidin-based SMLLM on actin-associated membrane fiber structures of DC2.4 cells. Phalloidin single-molecule labeling identified an average thickness of 52 nm for the actin-associated membrane fiber structures in DCs. Following LPS treatment, actin is redistributed from podosomes to cytoskeletal fibers. F-actin rearrangements upon LPS treatment can be transient.^56^ To that end, live superresolution imaging, i.e., using a complementary superresolution technique such as structured illumination microscopy (SIM), and live-cell-compatible probes such as SiR-Actin, live-cell SMLM techniques will offer new insights into their dynamics and related physiological functions.^58,59^ While phalloidin-based SMLLM confirmed the F-actin involvement in DC membrane fibers, the presence and spatial distribution of the major histocompatibility complex (MHC) molecules, for example, will provide additional insights related to the antigen presentation function of these membrane structures. To this end, SMLLM combining phalloidin and specific antibodies^44,60^ with optimized protocols may provide multiplexed capability.

Phalloidin-based SMLLM is relatively slow compared to PAINT imaging, with a typical imaging experiment having a frame rate of 30 frames *per* minute. In comparison, related techniques such as IRIS (frame rate: 20 Hz) and PAINT (frame rate: 25 Hz) using the actin probe Lifeact offer faster imaging speeds. While Lifeact has demonstrated faster imaging and high labeling density in PAINT and IRIS techniques, it has been shown to be less effective in labeling delicate structures.^27,35,36^ The binding of actin probes to F-actin is intricately governed by biochemical interactions dictated by the F-actin architecture, necessitating careful consideration of actin probes for specific investigations.^34^ Further studies to enhance the association and dissociation dynamics of protein-specific organic and protein-based labels are needed.

Quantitative characterization of phalloidin-AF647-F-actin interactions (on/off kinetics) in the presence and absence of KSCN would provide mechanistic insights into the chaotropic perturbation and the resultant increase in phalloidin-AF647 dissociation. However, affinity characterization of the phalloidin-AF647-F-actin interaction remains challenging. *In vitro* characterizations are challenging due to the polymeric nature of the substrate, F-actin. The use of fluorescence intensity decay analyses for *in situ* characterization of the dissociation rate^47^ is complicated because phalloidin cannot be cross-linked to the substrate *via* post-fixation (**Figure S9**). Alternatively, we have quantitatively evaluated the enhancement of the sampling rate in the presence of KSCN by fitting the cumulative number of binding events to a linear regression model. The cumulative number of single-molecule events *per* 1,000 frames fit into a linear regression model for acquisitions in the presence and absence of KSCN (**Figure 2h**). The presence of KSCN in the single-molecule labeling image acquisitions resulted in an approximately 3-fold increase in the sampling rate. While the reversibility of phalloidin-F-actin interaction due to promoted dissociation is a contributing factor, single-molecule labeling also relies on cumulative single-molecule labeling over time to achieve SMLM, particularly in the absence of KSCN.^44^

While phalloidin-staining is widely used in actin cytoskeletal superresolution imaging, the accelerated dissociation of phalloidin-AF647 reduces the labeling density for F-actin structures in *d*STORM experiments. While this phenomenon has a minimal impact on F-actin-abundant architectures, such as the cytoskeleton of adherent U2OS cells and podosomes of DC2.4 cells (**Figure 2e** and **3e**), it significantly affects the labeling density of membrane fiber structures (**Figure 3f-g**). Chemical cross-linking of the phalloidin-AF647 to F-actin is a plausible solution to prevent phalloidin dissociation. However, phalloidin-AF647 cannot be efficiently cross-linked to F-actin, i.e., using aldehyde-based chemical fixation (Figure S9).

F-actin-associated membrane structures are mechanically fragile, making them challenging to preserve. As phalloidin-based SMLLM does not require the step of phalloidin incubation and the consequent washing steps, it reduces the mechanical shear from additional washing steps. F-actin fixation followed by a short permeabilization step (methods) enabled the visualization of intracellular actin, proximal membrane fibers, and some distal membrane structures. An adapted fixation protocol optimized for TNT structures better preserved the distal membrane fiber network. ^15^ While optimized preservation protocols still need to be developed, phalloidin-based SMLLM presents an effective superresolution tool to study fragile membrane structures. In addition, the morphology of the membrane fibers is not affected by KSCN (Figure S10).

The quality of SMLM for life sciences studies depends on the labeling probes and preservation of the biological sample, particularly for nanostructures that can be easily damaged during sample preparation. Therefore, an important endeavor in advancing SMLM is to develop gentle labeling strategies that minimally perturb the biological sample. Phalloidin-based SMLLM demonstrates a new strategy to quantitatively super-resolve F-actin in the cell body and mechanically fragile cytoskeletal protrusions.

## METHODS

### Materials and reagents

DMEM (11960069-500ml), RPMI (11875093) penicillin-streptomycin (15140-122-100ml) L-Glutamine (25030-081-100ml), and 1x DPBS (14190-144-500ml) were purchased from Gibco. FBS (F0926-500ml), triton X-100 (9002-93-1-1L), MES (M3671-50G), EGTA (E3889-25G), sodium phosphate (342483-500G), potassium thiocyanate (207799-100G), DMSO (D8418-500ML), glucose oxidase from *Aspergillus niger* (G7141-50KU), catalase from bovine liver (C40-100MG), 2-mercaptoethanol (M6250-100ML), hydrochloric acid (258148-500ML) were ordered from Sigma-Aldrich. Alexa Fluor™ 647 Phalloidin (A22287), eBioscience™ Lipopolysaccharide (00497693), Cytiva HyClone™ HEPES Solution (SH3023701) and Cytiva HyClone™ Non-Essential Amino Acids 100X Solution (SH3023801) were purchased from Thermo Fisher Scientific. Potassium chloride (7447-40-7-500G) was obtained from Fisher Scientific. Paraformaldehyde (16%, 15710), and glutaraldehyde (10%, 16120) were purchased from Electron Microscopy Sciences. Magnesium chloride (7791-18-6-500G) was obtained from ACROS Organics. 8 well-chambered cover glasses (Sterile, No. 1, C8-1-N) were ordered from Cellvis. Unconjugated gold colloids (15711-20) were purchased from Ted Pella Inc.

The following materials were utilized for preparing and seeding bone marrow-derived dendritic cells (BMDCs). RPMI (SH30027.01) was purchased from Cytiva. 2-mercaptoethanol (31350010) was purchased from Gibco. FBS (45000-736) and penicillin-streptomycin 16777-164) were obtained from VWR. GM-CSF (576306) was obtained from BioLegend. A 100mm x 20mm petri dish (353003) was purchased from Corning.

### Buffers

Cytoskeleton buffer: MES (10 mM, pH 6.1), Potassium chloride (90 mM), Magnesium chloride (3 mM), and EGTA (2 mM). Fixation buffer 1: Paraformaldehyde (3.7%), Glutaraldehyde (0.1%), Triton-X-100 (0.5%) in Cytoskeleton buffer. Fixation Buffer 2: Paraformaldehyde (3.7%), Glutaraldehyde (0.1 %) in DPBS. Fixation Buffer 3: Paraformaldehyde (2%), Glutaraldehyde (0.05%), HEPES (0.2 M) in DPBS Fixation Buffer 4: Paraformaldehyde (4%), HEPES (0.2 M) in DPBS. Postfixation buffer: Paraformaldehyde (3.7%) in DPBS. Permeabilization Buffer: Triton-X-100 (0.1%) in DPBS. Buffer A: Tris (10 mM, pH 8.0), Sodium chloride (50 mM). Buffer B: Tris (10 mM, pH 8.0), Sodium chloride (50 mM), Glucose (10%). Glox solution: Glucose oxidase (14 mg), Catalase (17 mg/mL), in buffer A (200 μL). STORM buffer: Glox solution (7 μL), 2-Mercaptoethanol (7 μL), in buffer B (690 μL).

### Cell culture

U2OS cells (ATCC HTB-96) were cultured in DMEM supplemented with 10% FBS, 2 mM L-Glutamine, and 100 units/mL penicillin-streptomycin. DC 2.4 cells (Millipore SCC142) were cultured in RPMI supplemented with 10% FBS, 100 units/mL penicillin-streptomycin, 1x HEPES, and 1x NEAA. All cell lines were maintained at 37 °C in a humidified atmosphere of 5% CO_2_ and split at the confluence.

### Preparation of bone marrow-derived dendritic cells (BMDCs)

The bone marrow was harvested from the femurs of freshly euthanized BALB/c mice. The collection of bone marrow from mice was approved by the Animal Care Committee (ACC) at the University of Illinois Chicago (Protocol number 21-098). Bone marrow cells were then seeded onto a petri dish (100 mm x 20 mm) at a concentration of 2 × 10^5^ cells/mL and cultured in RPMI-1640 media supplemented with 10% FBS, 1% penicillin-streptomycin, 50 µM 2-mercaptoethanol, and 20 ng/mL GM-CSF. Cells were maintained at 37 °C in a humidified atmosphere of 5% CO_2_. Cell media was refreshed on days 3 and 6. Differentiated BMDCs were harvested on day 8 and collected in RPMI-1640 media supplemented with 10% FBS and 1% penicillin-streptomycin. Cells were centrifuged at 350 x g for 5 min.

### Sample preparation

#### Cell fixation

1) For imaging cytoskeletal actin in U2OS cells: U2OS cells were seeded (approximately 5,000 cells/well) in a chambered coverglass and grown in an incubator at 37 °C and 5% CO_2_. Following 36 h of incubation, cells were fixed and permeabilized in the freshly prepared fixation buffer 1 for 20 min at room temperature. Cells were maintained in PBS at 4 °C until imaging. 2) For imaging cytoskeletal actin and proximal membrane fibers in DC2.4 and BMDC cells: DC2.4 cells were seeded (approximately 2,500 cells/well) in a chambered coverglass and grown in an incubator at 37 °C and 5% CO_2_ for 24-48 h. BMDCs (approximately 5,000 cells/well) were seeded in a chambered coverglass and incubated for 16-24 h at 37 °C and 5% CO_2_. Following incubation, 3 drops of fixation buffer 2 were added into the medium of each well and incubated for 4 min at room temperature. All consequent solution exchange steps were performed with a Gilson PIPETMAN P200 pipette placed at the edge of the sample well. The chamber slides were held at an angle such that the solution slowly rose up, filling the well. The solution dispensing rate was approximately 400 µL/min. Performing solution exchanges at a slow rate is pivotal to minimizing damage to delicate structures. The media were replaced with fixation buffer 2 and incubated for 10 min at room temperature. The sample was then incubated with the permeabilization buffer for 5 min. Permeabilization buffer was replaced with DPBS, and the sample was imaged immediately after. 3) For imaging distal membrane fibers with extremely low actin contents, the following fixation procedure was adapted from Saida Albounit *et al*. ^26^ Briefly, 3 drops of freshly prepared fixation buffer 3 was added into the medium of each well and incubated for 4 min at 37 °C. All consequent solution exchange steps were performed with a Gilson PIPETMAN P200 pipette placed at the edge of the sample well. The chamber slides were held at an angle such that the solution slowly rose, filling the well. The solution dispensing rate was approximately 400 µL/min. Performing solution exchanges at a slow rate is pivotal to minimizing damage to delicate structures. The media containing drops of fixation buffer 3 was removed and replaced with fixation buffer 3. The chamber slide was incubated for 20 min at 37 °C and 5% CO_2_. Fixation buffer 3 was replaced by fixation buffer 4 and incubated for 20 min at room temperature. Fixation buffer 4 was replaced with DPBS for 30 s, and the sample was imaged immediately after. The sample chambers were incubated with a 1:5 dilution of gold nanoparticles to be used as fiducials.

#### LPS treatment

Overnight grown DC2.4 cells were treated with LPS-supplemented culture medium (5 μg/mL) for 21 h in an incubator at 37 °C and 5% CO_2_. The cells were fixed and permeabilized as previously described and imaged immediately afterward.

#### Phalloidin-AF647 handling

Phalloidin-AF647 was constituted in 150 μL of DMSO to a concentration of 66 μM according to the manufacturer guidelines. All consequent phalloidin-AF647 dilutions were made in DMSO, aliquoted, and stored at −20 °C to avoid repeated freeze-thaw cycles. The longevity of the diluted phalloidin-AF647 aliquots, especially for single-molecule imaging, can be prolonged by storing them in a sealed container with a desiccant. The final DMSO percentage (v/v) in the imaging buffer was 1-3%.

#### Phalloidin-based SMLLM

For phalloidin-AF647-single-molecule labeling, fixed and permeabilized U2OS and DC2.4 were imaged in an imaging buffer containing phalloidin-AF647 supplemented with KSCN in 1xDPBS. The chaotropic salt KSCN is a denaturant at higher concentrations. The concentration of the KSCN (300 mM) was chosen to be sufficiently high to promote dissociation but within the non-denaturing concentration range (200 – 400 mM). ^61^ The phalloidin-AF647 concentration was adjusted to maintain a sufficient single-molecule event density.

#### STORM imaging of DC2.4

Fixed and permeabilized DC2.4 cells were incubated with Phalloidin-AF647 (0.165 μM) in 1xDPBS overnight at 4 °C. The following day, the phalloidin solution was replaced with STORM buffer and imaged immediately after.

#### Phalloidin-based SMLLM after dSTORM imaging

Fixed and permeabilized U2OS cells were incubated with phalloidin-AF647 (0.33 μM) in 1xDPBS for 2 h at room temperature. The cells were washed three times with DPBS, and *d*STORM imaging was performed immediately after in the STORM buffer. Following the *d*STORM acquisition, the sample was washed on stage three times, and phalloidin-based SMLLM was performed.

#### Evaluation of phalloidin staining loss

Fixed and permeabilized DC2.4 cells were stained with phalloidin-AF647 (0.33 μM) for 2 h at room temperature. The cells were washed three times with DPBS. Fluorescent imaging was performed in 1xDPBS buffer.

#### Cross-linking of the phalloidin-staining

Fixed and permeabilized DC2.4 cells were stained with phalloidin-AF647 (0.33 μM) for 2 h at room temperature. The cells were washed three times with DPBS. The cells were then fixed in the post-fixation buffer for 15 min at room temperature. The cells were washed three times with DPBS, and fluorescence imaging was performed in the DPBS buffer.

#### SiR Actin Labeling

SiR actin constituted in DMSO (Stock concentration: 1 mM) was diluted in the DC growth medium to a final concentration of 1 µM. The culture medium of overnight grown DC cells on chambered cover glasses was replaced with the staining solution and incubated overnight (12h) in an incubator at 37 °C and 5% CO_2_.

### Microscopy

Fluorescence/Superresolution imaging was performed at a HILO/near TIRF setting on an inverted microscope (Nikon Instruments, Eclipse Ti2E). A 100×/1.49 oil-immersion objective (CFI Apochromat TIRF 100XC) was used with a 1.5x external magnifier. Single-molecule movies were recorded using a Prime 95B sCMOS camera at 16-bit with 2×2-pixel binning, creating an effective pixel size of 147 nm.

#### Fluorescence imaging

Fluorescence imaging of DC2.4 cells for the phalloidin-AF647 labeling loss experiments, after phalloidin-AF647 cross-linking and SiR actin labeling was performed using a 641 nm laser at an integration time of 50 ms and a laser power density of 2 W/cm^2^. Three consecutive images were captured, followed by captures at 40 min and 80 min, as intended. Fluorescence imaging of the DC2.4 cells for all other experiments was performed at an integration time of 50 ms and laser power density of 64 W/cm^2^.

#### *Phalloidin-*based SMLLM

A 641 nm laser was used at a power density of 251 W/cm^2^cm^2^ and a 2 s non-illuminating interval between consecutive frames. Due to the relatively slow kinetics of phalloidin, we used an integration time of 400 ms in the single-molecule labeling experiments to achieve an SNR of 1:3 or above. Single-molecule labeling of U2OS cells included an additional periodic photobleaching step at every 20^th^ frame at an integration time of 200 ms and a laser power density of 1.2 kW/cm^2^ to reduce out-of-focus fluorescence background. Single-molecule movies of 8,000-30,000 frames were recorded.

#### dSTORM imaging

*d*STORM imaging was performed under continuous illumination of 647 nm (1.046 kW/cm^2^). During the movie, a 405-nm laser was gradually increased from 8–32 W/cm^2^ to illuminate the sample and maintain a convenient density of activated molecules. *d*STORM movies of 20,000–80,000 frames were collected for DC2.4 and 100,000-120,000 for U2OS cells.

### Data processing

Superresolution image reconstruction was performed using the open-source ImageJ plug-in ThunderSTORM. ^62^ The camera settings were pixel size of 147 nm, photoelectrons per A/D count 0.76 (Prime 95B sCMOS camera at 16 bit, serial number: A21C203010), and base level 100. Drift correction was performed using fiducial beads that were present during the entire acquisition. Events with a sigma value greater than 170 were removed in the post-processing to reduce the diffusive appearance. All reconstructions are visualized at a pixel size of 20 nm in the reconstructed image. Results were visualized using either the ‘Normalized Gaussian’ with a forced uncertainty of 20 nm or the ‘Average Shifted Histogram’ with a lateral shift of 2 or 4.

The FFT analysis was performed by using the SciPy function, fft.fftn, in Python (version 3.12.1). Cross-sectional intensity profiles along the membrane fibers for each technique (n = 9) were obtained in ImageJ. Their corresponding FFTs were calculated, keeping only the positive frequencies. The SciPy function, find_peaks, was then used to identify peaks to display the frequency components of the *d*STORM and phalloidin-based SMLLM methods in frequency space. Equal threshold values were used for peak finding in the FFT.

The analysis of the actin densities in DC cells was conducted using ImageJ (version 1.54f). The distinct regions of interest (ROIs) were selected based on the intensity of the image using the ‘threshold’ option under the ‘adjust’ menu. Subsequently, distinct actin architectures were further isolated based on the size and the circularity using the ‘analyze particles’ tool in the ‘analyze’ menu (Table S1). The ‘area’ and ‘RawIntDen’ of the ROIs were measured using the ‘measure’ command in the ‘analyze’ menu. ‘RawIntDen’ is the sum of the pixel values in a selected ROI. The actin density *per* unit area was obtained by dividing the ‘RawIntDen’ value by the area of an ROI.

### Statistical information

Statistical significance was evaluated using an unpaired Student’s *t-test*, two-tailed.

## ASSOCIATED CONTENT

### Supporting Information

The following files are available free of charge.

Chosen ROIs used in quantitative analysis of F-actin densities; Phalloidin-based SMLLM revealed distal membrane fibers with low actin contents; SiR-Actin labeling of live DC2.4 cells reveals low-actin contents in distal membrane fibers compared to the cell body; Increasing *d*STORM acquisition times did not improve the labeling density of membrane fibers; Comparison of the labeling consistency between *d*STORM and phalloidin-based SMLLM in the frequency space; Phalloidin-based SMLLM reveals nanoscale F-actin rearrangements upon LPS treatment of DC2.4 cells; LPS-DC2.4 cells displayed a defined cytoskeleton; The selected ROIs for actin density calculations; Chemical cross-linking does not affect phalloidin-AF647 dissociation in the STORM imaging buffer; KSCN does not impact membrane fiber structures. ImageJ parameters used in the DC2.4 actin density analysis.

## AUTHOR INFORMATION

### Corresponding Author

Ying S. Hu – Department of Chemistry, College of Liberal Arts and Sciences, University of Illinois Chicago, Chicago, IL, 60607-7061, USA; orcid.org/0000-0002-3912-7063;

### Author Contributions

HG and YSH conceived and planned the experiments, HG performed single-molecule labeling imaging experiments and data analyses, HG and TP performed STORM imaging, TP, and JA assisted in data analyses, JB performed FFT analysis, C-J.C. and ZZ prepared BMDC, BS, and TP prepared BMDC and DC2.4 samples for imaging, and HG and YSH wrote the manuscript.

### Data availability

All data will be available upon request to the corresponding author.

## ACKNOWLEDGMENT

The authors would like to thank Dr. Carlos Murga-Zamalloa for the gift of Phalloidin-647, Dr. Ruixuan Gao for his assistance with confocal microscopy, and Dr. Yu-Shiuan Cheng for his support with phalloidin handling.

