## Supplementary material for "Quantitative Superresolution Imaging of F-Actin in the Cell Body and Cytoskeletal Protrusions Using Phalloidin-Based Single-Molecule Labeling and Localization Microscopy": Figure S1-S10, Table S1

This PDF file includes: Figures S1 to S10 and Table S1


| | |
|---|---|
| **Figure S1** | Chosen ROIs used in quantitative analysis of F-actin densities. |
| **Figure S2** | Phalloidin-based SMLLM revealed distal membrane fibers with low actin contents. |
| **Figure S3** | SiR-Actin labeling of live DC2.4 cells reveals low-actin contents in distal membrane fibers compared to the cell body. |
| **Figure S4** | Increasing *d*STORM acquisition times did not improve the labeling density of membrane fibers. |
| **Figure S5** | Comparison of the labeling consistency between *d*STORM and phalloidin-based SMLLM in the frequency space. |
| **Figure S6** | Phalloidin-based SMLLM reveals nanoscale F-actin rearrangements upon LPS treatment of DC2.4 cells. |

| | |
|---|---|
| **Figure S7** | LPS-DC2.4 cells displayed a defined cytoskeleton. |
| **Figure S8** | The selected ROIs for actin density calculations. |
| **Figure S9** | Chemical cross-linking does not affect phalloidin-AF647 dissociation in the STORM imaging buffer. |
| **Figure S10** | KSCN does not impact membrane fiber structures. |
| **Table S1** | ImageJ parameters used in the actin density analysis |

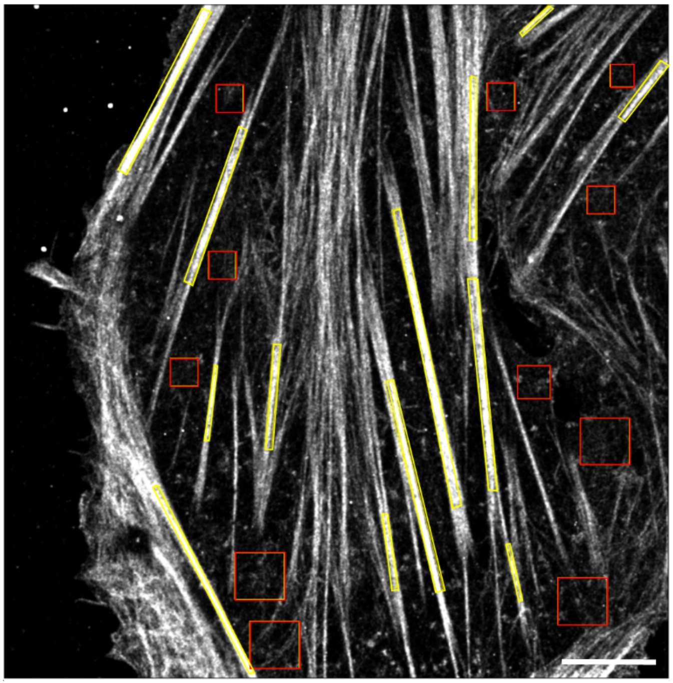

**Figure S1. Chosen ROIs used in quantitative analysis of F-actin densities.** Stress fibers: yellow boxes; Thin fibers: Red boxes. Scale bar: 5 μm

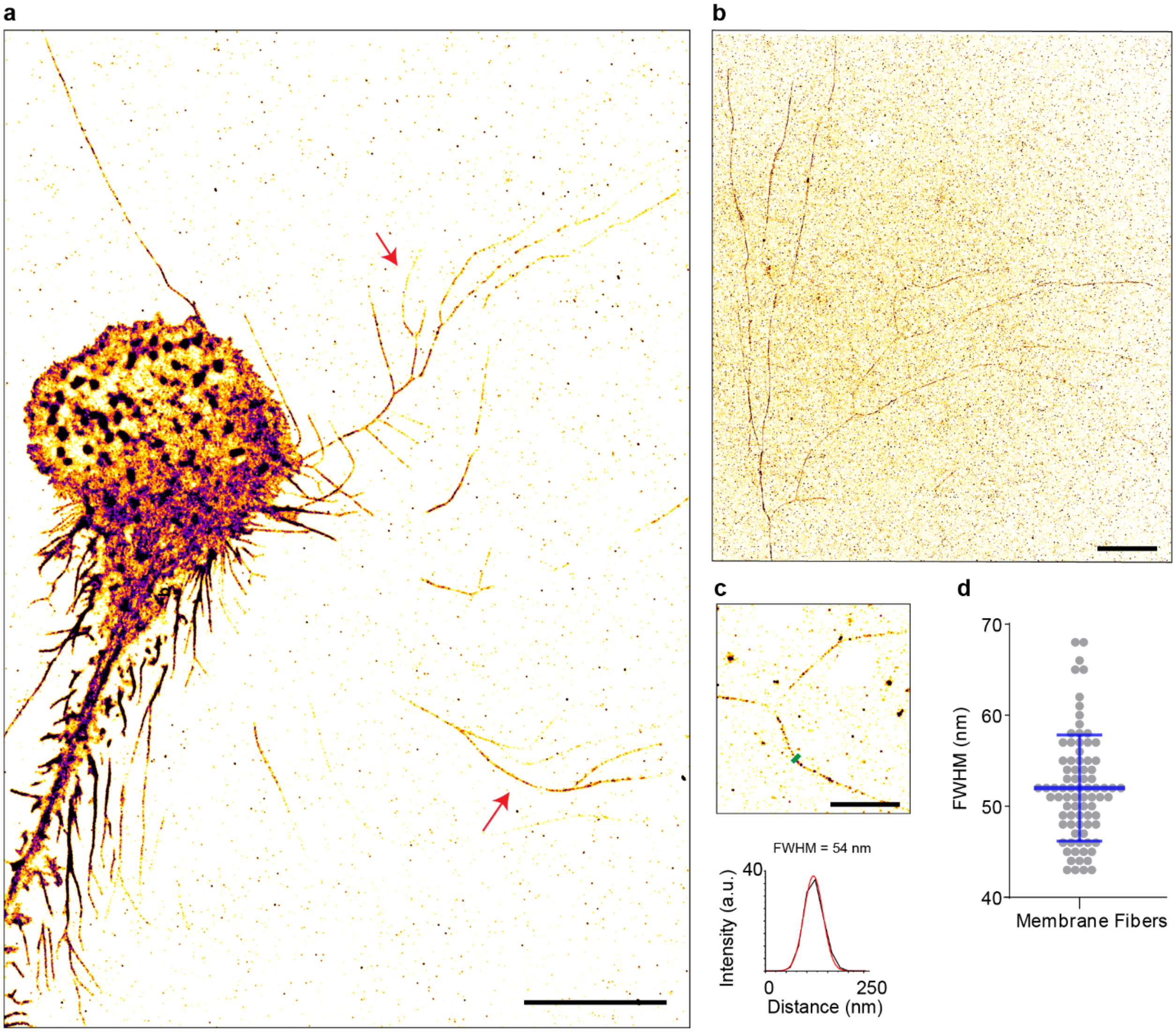


**Figure S2. Phalloidin-based SMLLM revealed distal membrane fibers with low actin contents. a.** Phalloidin-based SMLLM image of a DC2.4 cell exhibiting extending membrane fiber structures (red arrows). **b.** Actin-associated membrane fiber structures extending further from the cell body (within the TIRF field-of-view of approximately 80×80 µm). **c.** A representative zoomed-in view of the extended membrane fiber network (top), and the cross-sectional intensity profiles along the line indicated in green (bottom). The Gaussian fitting of the average intensity profile is shown in red. **d.** Distribution of FWHM along membrane fiber structures from two independent experiments. ($n = 82$). Scale bars: 10 µm (**a** and **b**), and 2 µm (**c**).

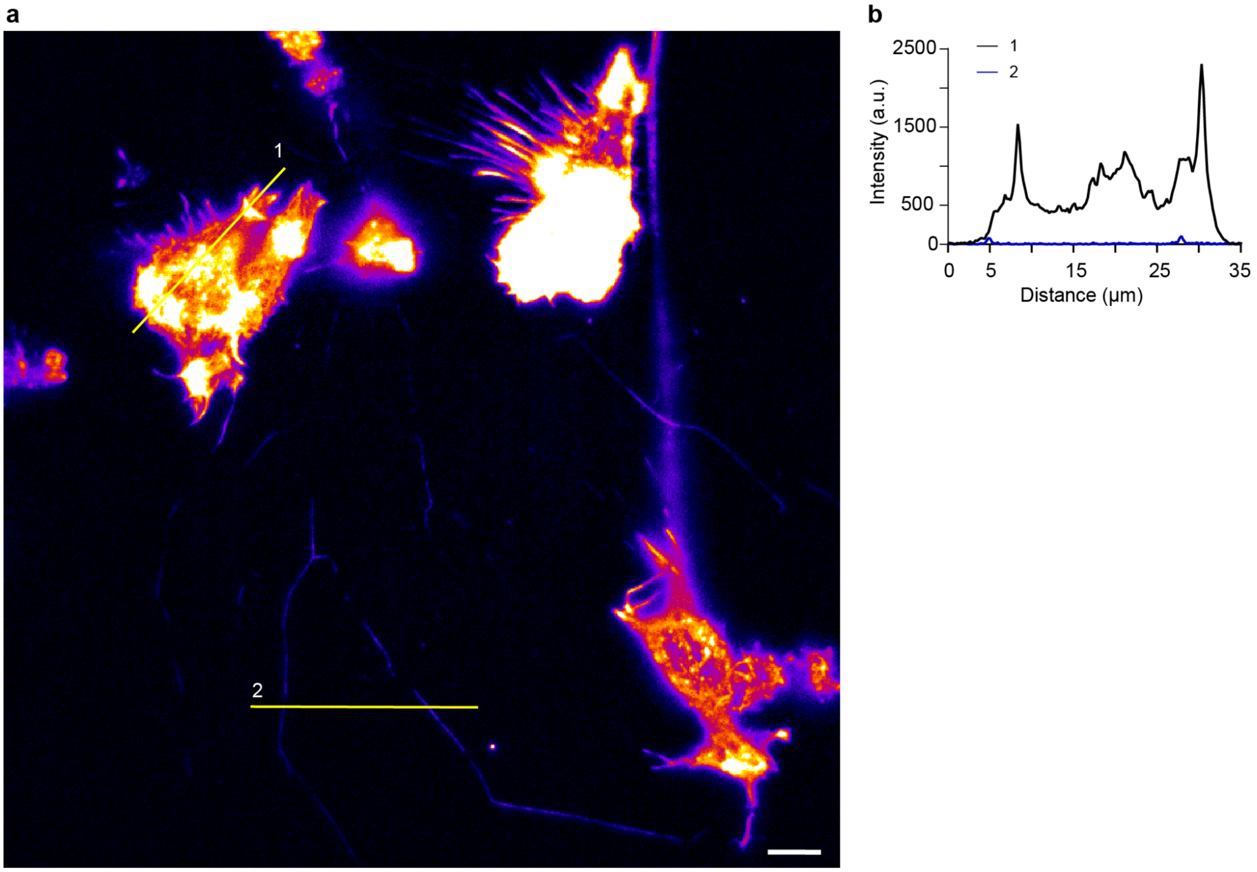


**Figure S3. SiR-Actin labeling of live DC2.4 cells reveals low-actin contents in distal membrane fibers compared to the cell body. a.** Fluorescence image of SiR-actin-labeled live DC2.4 cells. **b.** Representative cross-sectional line profiles through a DC2.4 cell and membrane fibers. Scale bar: 10 µm

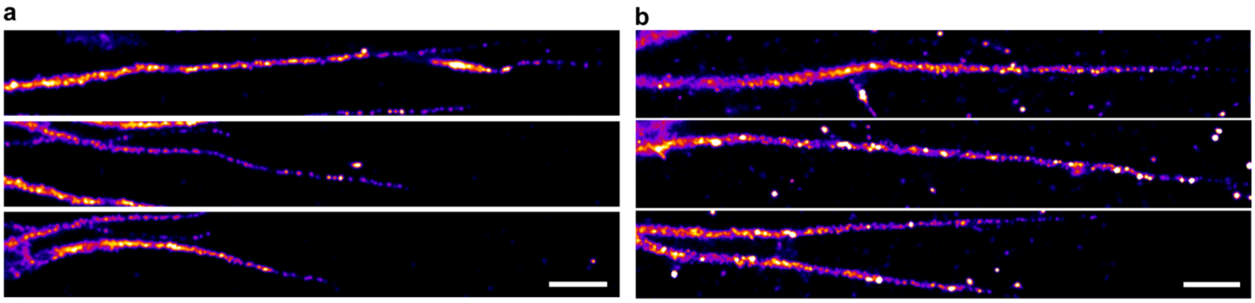

**Figure S4. Increasing *d*STORM acquisition times did not improve the labeling density of membrane fibers. a.** Collage of membrane fibers from a *d*STORM acquisition reconstructed from ~30000 frames. **b.** Collage of membrane fibers from a *d*STORM acquisition reconstructed from ~78000 frames. Scale bars: 1 µm

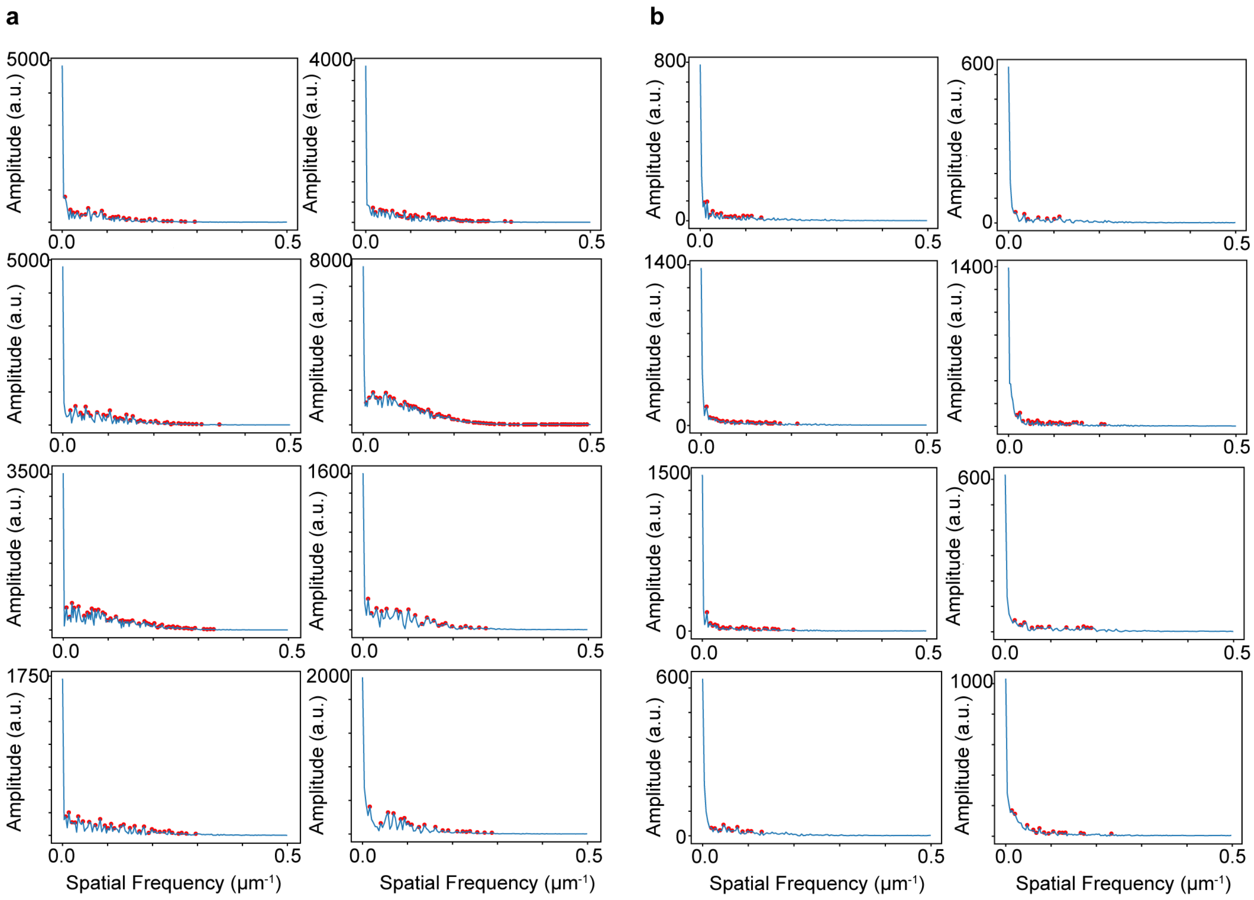


**Figure S5. Comparison of the labeling consistency between *d*STORM and phalloidin-based SMLLM in the frequency space. a.** Representative FFT plots of cross-sectional intensity profiles along eight membrane fibers reconstructed from three independent *d*STORM experiments. The red circles highlight detected frequency components. **b.** Representative FFT plots of cross-sectional intensity profiles along eight membrane fibers reconstructed from three independent phalloidin-based SMLLM experiments. The red circles highlight detected frequency components.

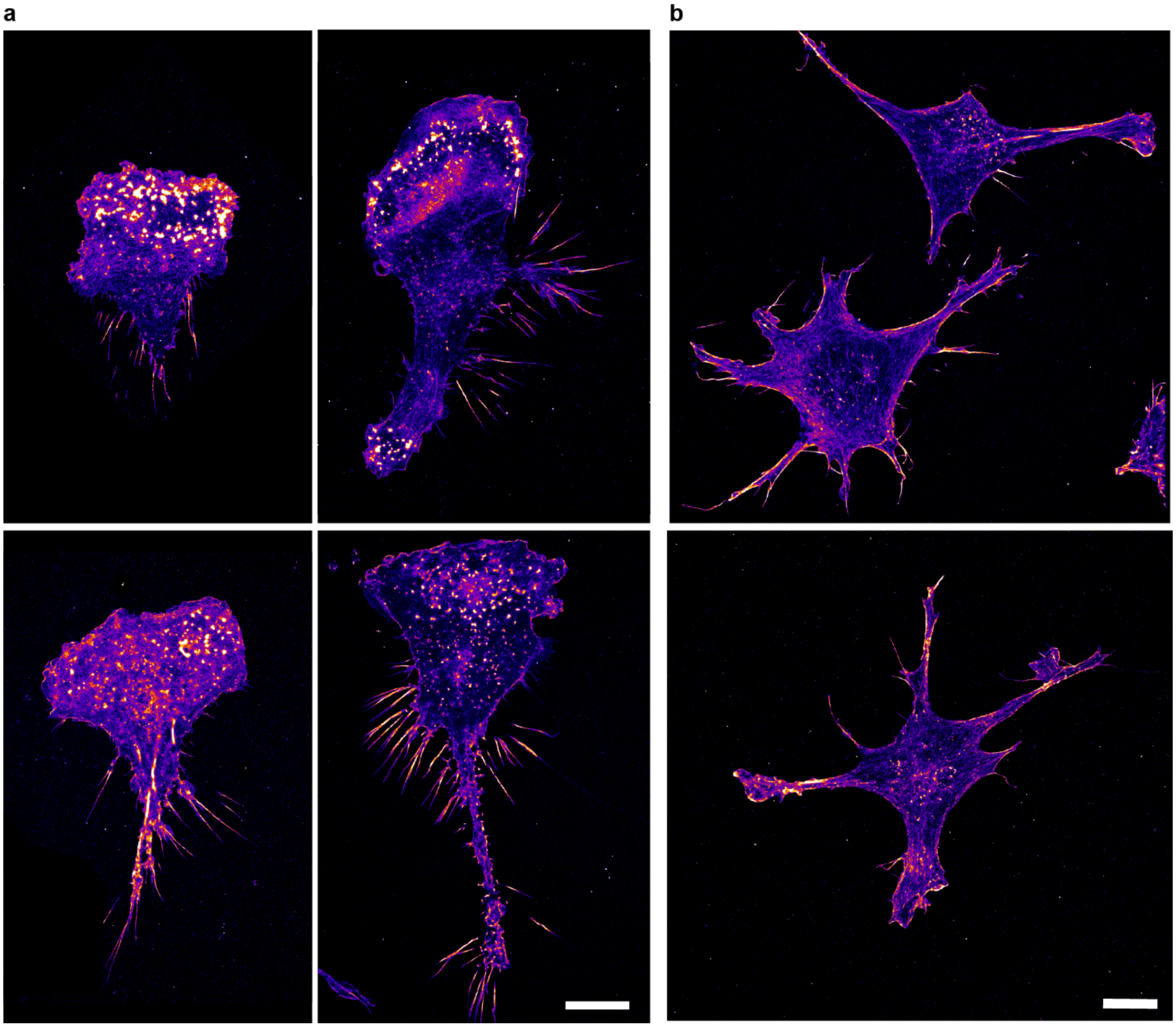


**Figure S6. Phalloidin-based SMLLM reveals nanoscale F-actin rearrangements upon LPS treatment of DC2.4 cells. a.** Phalloidin-based SMLLM image of iDC2.4 cells. **c.** Phalloidin-based SMLLM image of LPS-DC2.4 cells.

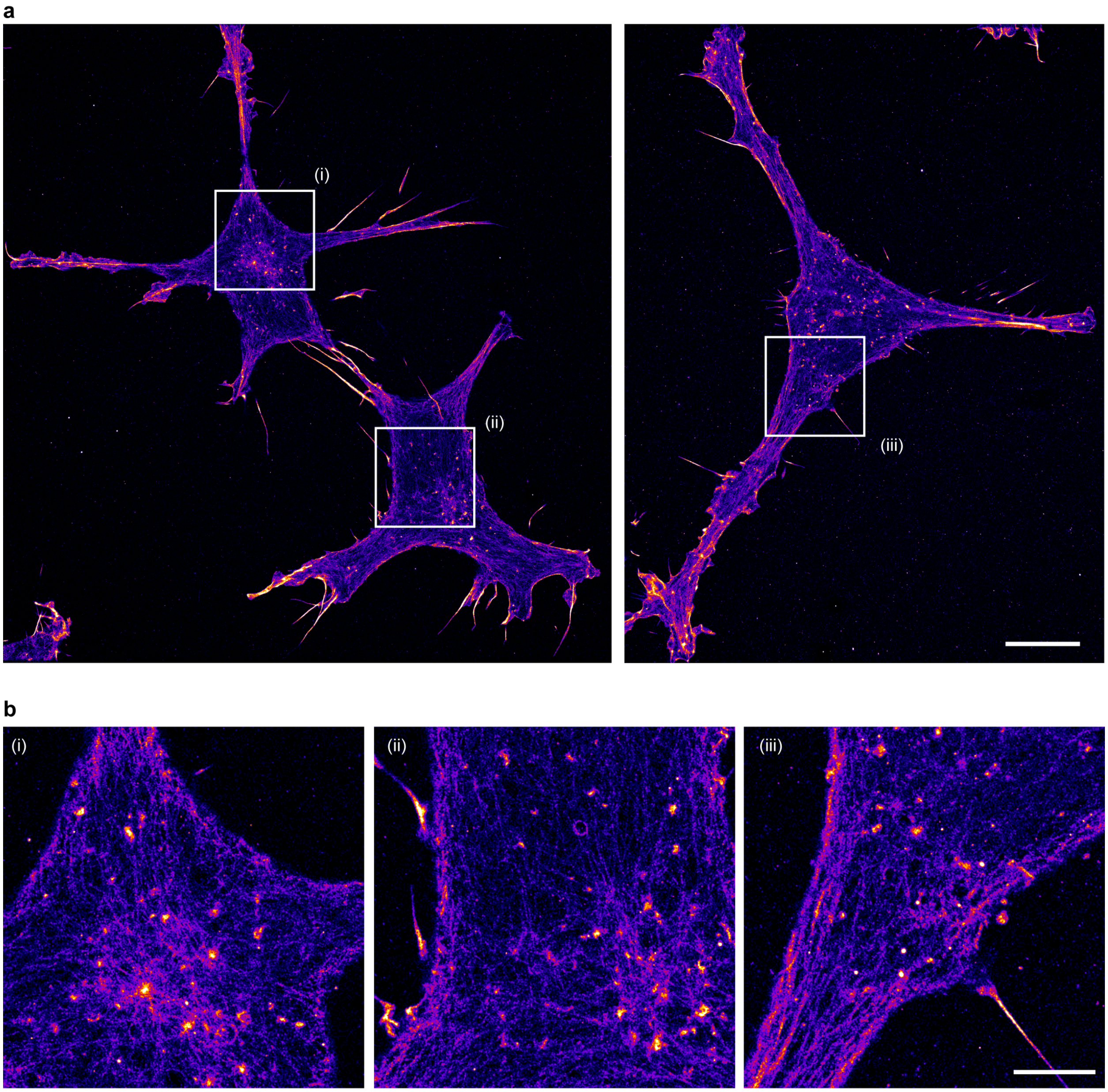


**Figure S7. LPS-DC2.4 cells displayed a defined cytoskeleton. a.** Phalloidin-based SMLLM images of LPS-DC2.4 cells. **b.** Zoomed in regions indicated in panel a. Scale bars: 10 μm (**a**), and 3 μm (**b**).

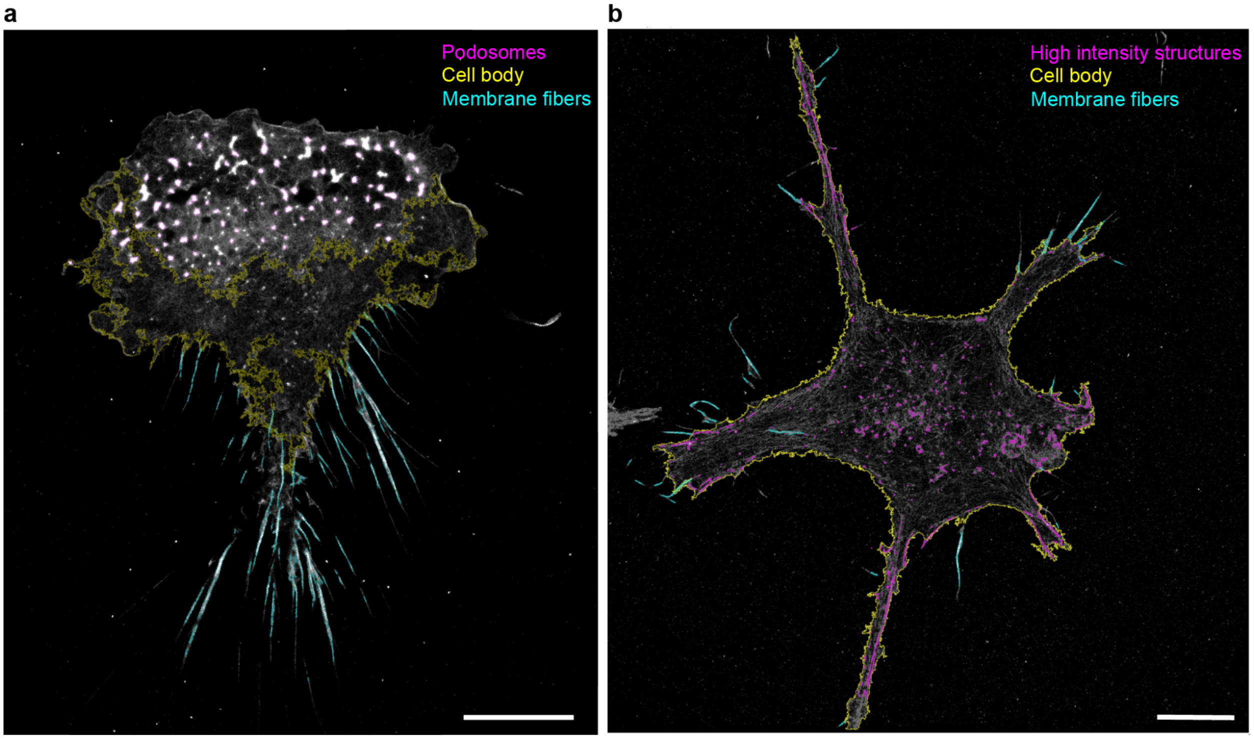


**Figure S8. The selected ROIs for actin density calculations. a.** iDC2.4 cell corresponding to Figure 4b. **b.** LPS-DC2.4 cell corresponding to Figure 4c. Scale bars: 10 µm

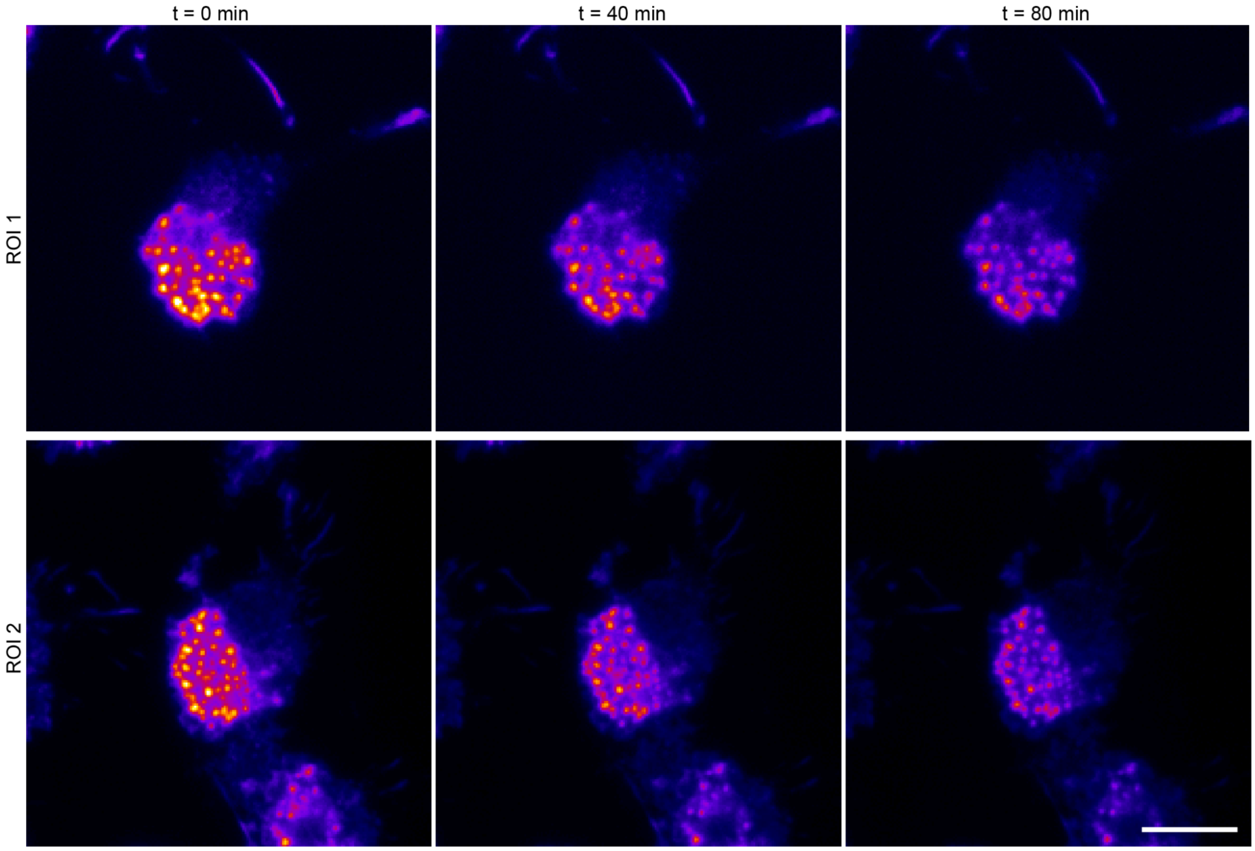


**Figure S9. Chemical cross-linking does not affect phalloidin-AF647 dissociation in the STORM imaging buffer.** The DC2.4 cells were stained with phalloidin-AF647 (2h, room temperature) and post-fixed. Fluorescence images of DC2.4 were taken at 40-minute time intervals in the STORM buffer. Scale bar: 10 μm

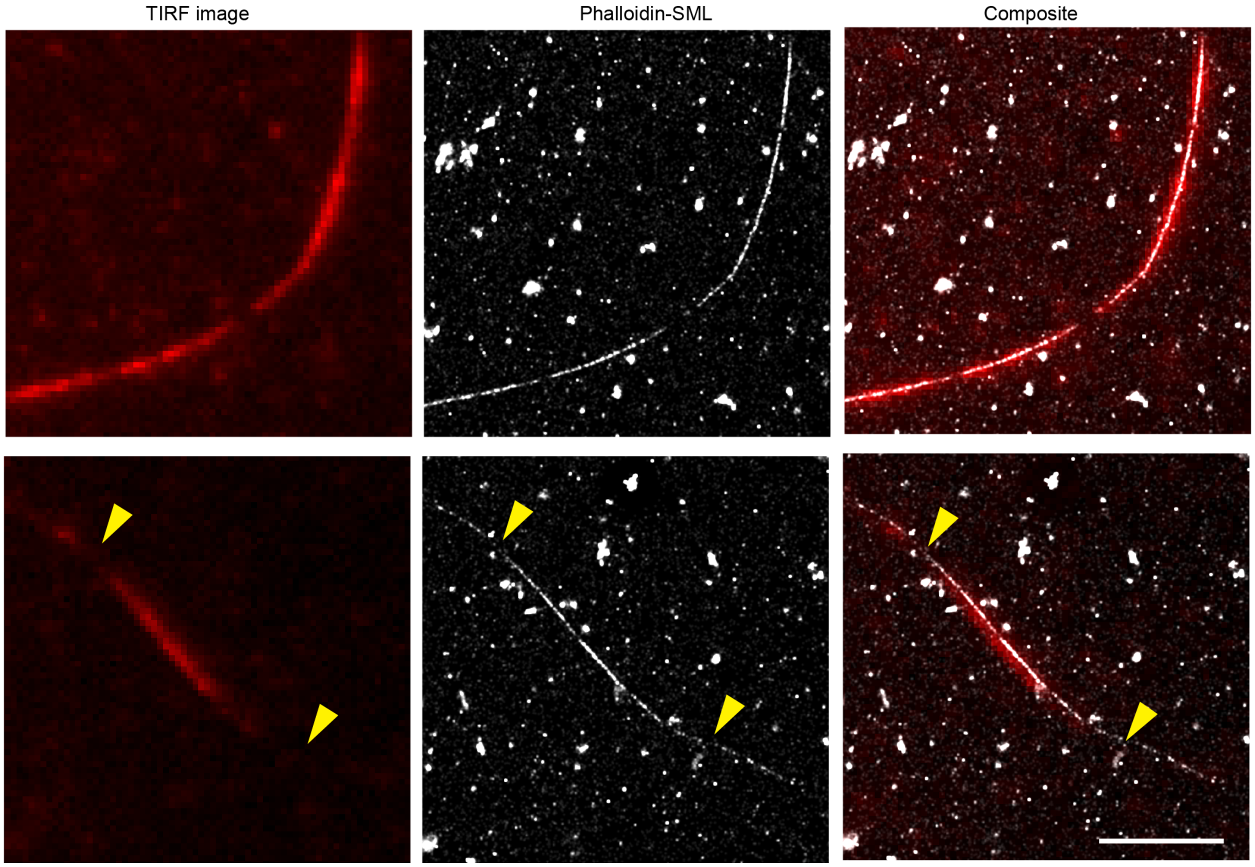


**Figure S10. KSCN does not impact membrane fiber structures.** Representative TIRF images (left), phalloidin-based SMLLM images (middle), and their composite images (right), taken on the same region of membrane fibers. Two representative regions of interest are shown (top and bottom). The membrane fibers were labeled with phalloidin-AF647 (165 nM, 30 min), and phalloidin-based SMLLM acquisitions were performed immediately after with no washing step. Arrowheads point to additional fiber regions captured by phalloidin-based SMLLM. Scale bar: 5 µm

**Table S1. ImageJ (version 1.54f) parameters used in the actin density analysis**

| Cell | ROI | Intensity threshold (a.u) | | Size (µm²) | Circularity |
|---|---|---|---|---|---|
| | | Min | Max | | |
| iDC2.4 | Podosomes | 9.73 | Default | 0.04-infinity | 0.4-1 |
| | Cell body | 0.41 | 1.63 | 0.04-infinity | 0-1 |
| | Membrane fibers | 4 | Default | 0.04-infinity | 0-0.3 |
| LPS-DC2.4 | Actin-rich structures[1] | 4 | Default | 0.02-infinity | 0.5-1 |
| | Membrane fibers[2] | 4 | Default | 0.04-infinity | 0-0.5 |
| | Cell body[3] | 0.88 | 1.59 | 0.04-infinity | 0-1 |

[1.]Selected membrane fibers were omitted before taking the measurements. [2.]Selected intracellular ROIs were omitted. [3.]The ROIs of actin-rich structures were deleted before taking the measurements. The cell was contoured, and measurements were taken.
